# An improved ChEC-seq method accurately maps the genome-wide binding of transcription coactivators and sequence-specific transcription factors

**DOI:** 10.1101/2021.02.12.430999

**Authors:** Rafal Donczew, Amélia Lalou, Didier Devys, Laszlo Tora, Steven Hahn

## Abstract

Mittal and colleagues have raised questions about mapping transcription factor locations on DNA using the MNase-based ChEC-seq method (Mittal et al., 2021). Partly due to this concern, we modified the experimental conditions of the MNase cleavage step and subsequent computational analyses, resulting in more stringent conditions for mapping protein-DNA interactions (Donczew et al., 2020). The revised method (dx.doi.org/10.17504/protocols.io.bizgkf3w) answers questions raised by Mittal et al. and, without changing earlier conclusions, identified widespread promoter binding of the transcription coactivators TFIID and SAGA at active genes. The revised method is also suitable for accurately mapping the genome-wide locations of DNA sequence-specific transcription factors.

---

ChEC-seq and other nuclease-based methods such as Cut&Run map protein locations on DNA by targeting nuclease activity to specific transcription factors and mapping DNA cleavages (Schmid et al., 2004; Skene and Henikoff, 2017; Zentner et al., 2015). For ChEC-seq, yeast cells expressing a protein-micrococcal nuclease (MNase) fusion are permeabilized, MNase is activated by the addition of calcium, and the resulting DNA fragments are mapped. This approach was used to map the genome-wide locations of transcription factors that do not directly bind DNA as well as sequence-specific DNA binding factors e.g., (Baptista et al., 2017; Grünberg and Zentner, 2017; Zentner et al., 2015). Potential advantages of this approach include avoiding non-specific protein-DNA crosslinking in highly transcribed regions and efficient mapping of factors that do not directly bind DNA.

In the original ChEC-seq method, DNA binding sites are mapped by identifying DNA cleavage frequencies at least 10-fold over the genome-wide average. The free MNase control (MNase with a nuclear import signal) was used for qualitative rather than quantitative comparisons (Zentner et al., 2015). Short times of digestion after calcium addition were used to maximize specific vs free MNase cleavages and the binding sites for some sequence-specific transcription factors were identified using ChEC-seq. However, Mittal et al. suggested that many binding locations identified by this approach, especially for factors that do not directly bind DNA, may be due to non-specific DNA cleavage. Partly due to this concern, we developed an improved ChEC-seq approach.

An important criterion is that binding sites identified by ChEC-seq should have DNA cleavage signals significantly stronger than the free MNase control. However, MNase cleavage activity is biased by the local chromatin environment and DNA sequence. Due to this property, data for different factors generated by ChEC-seq carry some qualitative resemblance to free MNase and to each other and thus, the quantitative differences in local DNA cleavage frequency identify specific versus non-specific DNA interactions.

To minimize non-specific DNA cleavage and to avoid over digestion at authentic binding sites, we modified the MNase cleavage conditions by using 10-fold lower calcium concentrations (0.2 mM final) and limited MNase digestion time to a single time point (5 minutes). We found that the 5 minute digestion at lower calcium concentration results in cleavage kinetics similar to that recommended by Zentner and colleagues with 2 mM final calcium concentration and incubation for 30 seconds (Zentner et al, 2021). With these conditions, the use of a fixed 5 minute time point allows many samples to be efficiently analyzed during a single experiment. Spike-in DNA is used to normalize samples for quantitative analysis. ChEC DNA cleavage patterns are compared with MNase controls where free MNase, with a nuclear import signal, is expressed from a promoter with greater or equal activity as the factor under study. Methods and the criteria for peak calling as well as the ChEC-seq results for SAGA and TFIID binding are described in (Donczew et al., 2020). A detailed protocol is available at protocols.io (dx.doi.org/10.17504/protocols.io.bizgkf3w).

With these improved ChEC-seq conditions and data analysis pipeline, ChEC-seq was performed for TFIID subunits Taf1, Taf7 and Taf13 (four biological replicates each). We identified >2900 binding sites for each TFIID subunit with >86% overlap between binding sites identified for each Taf. The average Taf-ChEC cleavage signals were far above that from free MNase and overlapped with many of the reported Taf1 ChIP-exo signals. By applying our computational criteria to published ChIP-exo data (Vinayachandran et al., 2018), we found ~75% overlap between TFIID binding sites mapped using ChEC-seq and ChIP-exo (Donczew et al., 2020).

We also used this approach to revisit genome-wide binding of the coactivator SAGA using Spt7-MNase (a SAGA-specific subunit) and identified >3500 promoter binding sites (Donczew et al., 2020). As with TFIID ChEC-seq, the average Spt7-ChEC signals were far above that observed for free MNase, a conclusion similar to the one reported by Bruzzone and colleagues for Gcn5 ChEC-seq (Bruzzone et al., 2021). We found that the binding of TFIID and SAGA does not discriminate against either of the two gene classes defined as: TFIID-dependent and coactivator redundant (CR) (Donczew et al., 2020). Importantly, our analysis revealed ~90% overlap between Taf7 and Spt7 binding, supporting our prior conclusions that most genes are regulated by both TFIID and SAGA (Baptista et al., 2017; Warfield et al., 2017). Our new results confirm that both these coactivators have extensive genome-wide promoter binding to active genes with little or no preference for different gene classes and clearly answer the concerns raised by Mittal et al.

Finally, to demonstrate the general utility of our approach using a different type of factor and to compare results with published ChIP-exo data, we mapped the binding of two yeast sequence-specific factors with different numbers of genome-wide binding sites: Abf1 and Rap1 (**Fig 1**). We found 1060 (Rap1) and 2308 (Abf1) bound promoters genome-wide, based on two independent experiments for each factor. Although ChEC-seq finds ~3-fold more Rap1 binding sites compared with ChIP-exo (Rhee and Pugh, 2011), ~80% of the ChIP-exo mapped binding sites are also identified using ChEC-seq. *De novo* motif discovery using 100 nt sequences surrounding peak summits found the Rap1 consensus binding site in the vicinity of the peaks for 43.4% and 56.7% of Rap1-bound promoters identified by ChEC-seq and ChIP-exo, respectively. From Abf1 mapping data, we found the known consensus binding site close to the peak summit in 971 out of 2308 bound promoters (42.1%) which further confirms reliability of modified ChEC-seq in mapping transcription factors. In addition, the number of promoters bound by Abf1 and Rap1 is very similar to ChEC-seq results obtained by Zentner and colleagues using different experimental conditions and data analysis approaches (Zentner et al., 2021).

**Figure 1.**
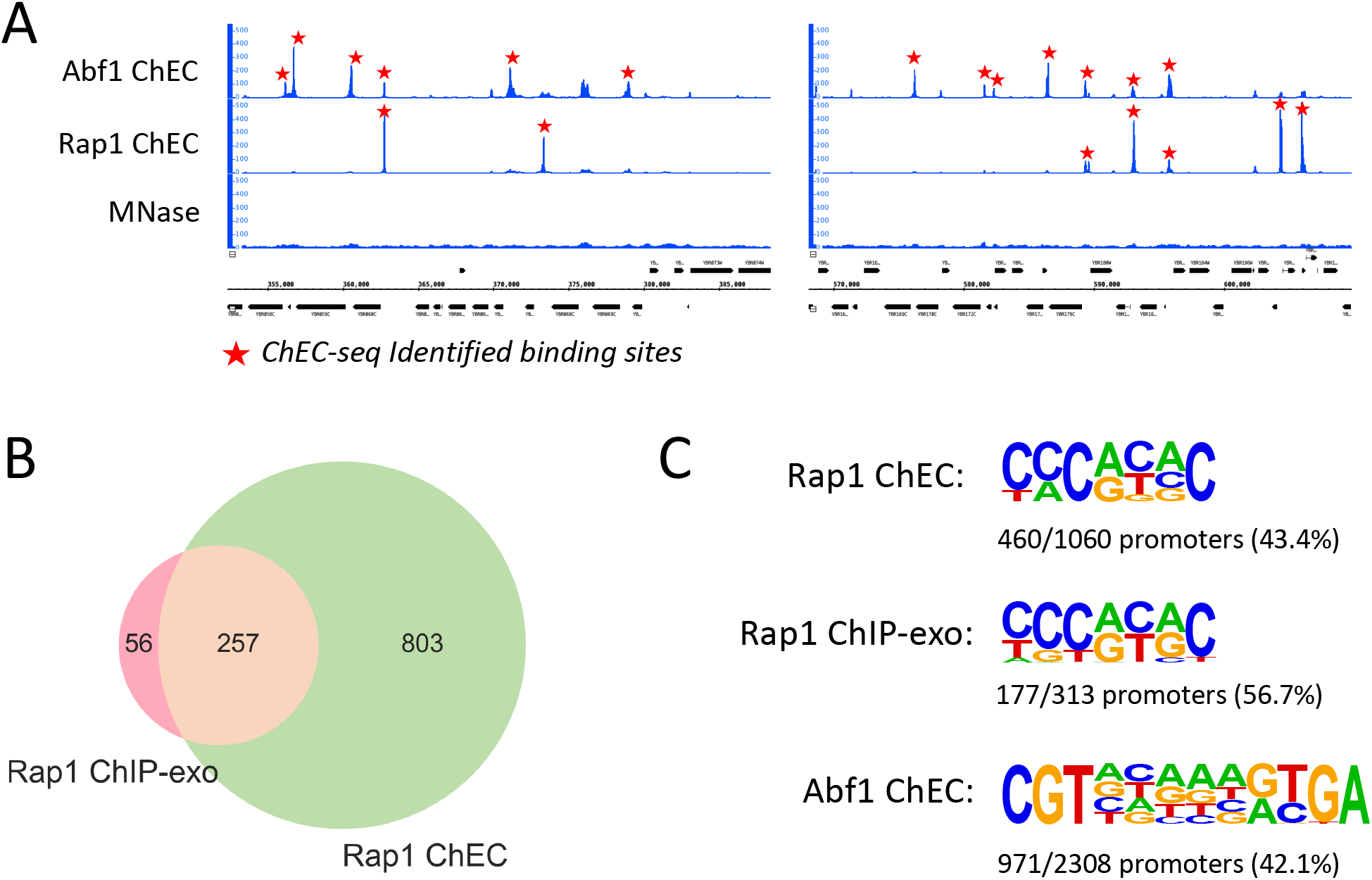
Mapping two yeast transcription factors with different genome-wide distributions. **(A)** DNA cleavage signals from Abf1 and Rap1 ChEC-seq at representative genomic locations compared to free MNase signal. Identified binding sites, found in both replicates, are marked by red stars. **(B)** Venn diagram showing the overlap of Rap1 bound promoters identified by ChEC-seq and ChIP-exo (Rhee and Pugh, 2011). **(C)** Sequence logos of the motifs enriched in a ± 50 bp window around promoter located peaks for Abf1 and Rap1 ChEC and Rap1 ChIP-exo experiments. The number and proportion of promoters showing the motif are shown.

In conclusion, the requirement of higher ChEC signals for factor-directed DNA cleavage compared with free MNase is an important criterion to use for identification of specific DNA binding sites. Results from our new experimental approach and computational analysis clearly meet this threshold for mapping the binding of the transcription coactivators TFIID and SAGA and for sequence-specific transcription factors.

## Data availability

ChEC-seq data is available at GEO (GSE142120).

## Author contributions

RD and AL developed the modified ChEC method and performed ChEC, RD performed all computational analysis, RD and SH designed the study, analyzed the data, and wrote the manuscript and all other authors edited and approved the manuscript.

## Acknowledgements

We thank Olivia Sommers for the Rap1 and Abf1 ChEC strains, Gabe Zentner, Steve Henikoff and David Shore for discussions and communication of unpublished results, and Frank Pugh and colleagues for communication of their concerns regarding ChEC-seq. Supported by NIH grants GM053451 and GM075114 to SH, by Agence Nationale de la Recherche (ANR)-18-CE12-0026 grant to D.D. and grant ANR-10-LABX-0030-INRT, a French State fund managed by the ANR under the frame program Investissements d’Avenir ANR-10-IDEX-0002-02 to L.T. and NIH P30 CA015704 to the FredHutch genomics and computational shared resources.

